# Reovirus sensitizes microsatellite stable colorectal cancer to anti-PD-1 treatment via cross-talk in innate and adaptive immune systems

**DOI:** 10.1101/2021.09.03.458915

**Authors:** Titto Augustine, Peter John, Tyler Friedman, Jeeshan Jiffry, Hillary Guzik, Rifat Mannan, Riya Gupta, Catherine Delano, John M. Mariadason, Xingxing Zang, Radhashree Maitra, Sanjay Goel

**Affiliations:** Department of Medicine, Albert Einstein College of Medicine, Bronx, New York; Department of Microbiology and Immunology, Albert Einstein College of Medicine, Bronx, New York; Department of Neuroscience, Florida State University, Tallahassee, Florida; Analytical Imaging Facility, Albert Einstein College of Medicine, Bronx, New York; Department of Pathology, City of Hope, Duarte, California; Columbia University, New York, New York; Gastrointestinal Cancers Program and Oncogenic Transcription Laboratory, Olivia Newton-John Cancer Research Institute, La Trobe University School of Cancer Medicine, Melbourne, Australia; Department of Urology, Albert Einstein College of Medicine, Bronx, New York; Department of Medical Oncology, Montefiore Medical Center, Bronx, New York; Department of Biology, Yeshiva University, New York, New York

**Keywords:** Colorectal cancer, translational, combinatorial therapy, reovirus, anti-PD-1, immune checkpoint, microsatellite instability

## Abstract

**Background:** Microsatellite stable (MSS) colorectal cancer (CRC) represents ^~^85% of all CRCs. These tumors are poorly immunogenic and largely resistant to immunotherapy, necessitating a need to develop new immune enhancing strategies. Oncolytic reovirus has a high propensity to replicate in *KRAS* mutant tumors which account for ^~^50% of MSS CRCs. Current study explores the ability of reovirus to potentiate the effect of immune checkpoint inhibition in MSS CRC.

**Methods:** Effectiveness of reovirus infection was quantified through MTT assay for cell viability, and expression of immune-response genes by flow cytometry, RT-qPCR, and microarray. Computational analysis of differentially expressed genes was performed by TAC, DAVID and STRING. Combinatorial approach using anti-PD-1 monoclonal antibody was assessed in *ex vivo* and *in vivo* models. Live-cell imaging, tumor volume and survival were measured for quantification of anti-tumor activity. Expression of pattern recognition receptors (PRRs), cell surface and activation markers of immune cells, and PD-1/PD-L1 axis were studied using multi-color flow cytometry, immunoblotting, immunohistochemistry, and immunofluorescence.

**Results:** Reovirus infection exerted growth arrest and expression of immune-response genes in CRCs cell lines in a KRAS-dependent manner. However, microsatellite instability, rather than *KRAS* status determined immune-repose pathways, functionalities and biological processes post-reovirus infection. Furthermore, reovirus significantly enhanced the anti-tumor activity of anti-human PD-1 [nivolumab] treatment in MSS CRC cell lines *ex vivo*. Similarly, reovirus increased the activity of anti-mouse PD-1 treatment in the CT26 [MSS, KRAS^Mut^], but not the MC38 [MSI, KRAS^Wt^] syngeneic mouse model of CRC. Combinatorial treatment has reduced the proliferative index, increased apoptosis and differentially altered PD-L1/PD-1 signaling among CT26 and MC38 tumors. Activation of innate immune system and expression of PRRs and antigen presentation markers were observed under reovirus and anti-PD-1 treatment that additionally reduced immunosuppressive macrophages. This led to an increase in T cell subsets, increase in effector T cell activation, and decrease in exhaustion markers specifically within CT26 microenvironment.

**Conclusion:** The current study systematically evaluates immune characteristics and immune microenvironment of CRC under reovirus/anti-PD-1 combination treatment that proves increased effectiveness among MSS compared to MSI CRCs. This is a promising regimen warranting translation into clinical trials.

**One Sentence Summary:** Oncolytic reovirus alters innate and adaptive immune system and potentiates MSS type colorectal cancer to checkpoint inhibition therapy.

## Background

Colorectal cancers (CRCs) with microsatellite instability (MSI) generate high levels of neoantigens due to mutations in DNA repair genes. The resulting neoantigens are detected by tumor infiltrating lymphocytes (TILs).^1^ Consequently, patients with MSI CRCs have experienced significant clinical benefit from immune checkpoint inhibition.^2^ Most of advanced MSI cancers express high levels of immune checkpoint proteins, including PD-1, PD-L1, CTLA-4, LAG-3, IDO, that help these cells evade immune destruction by TILs, and create an immunosuppressive tumor microenvironment (TME).^3 4^ Monoclonal antibodies targeting PD-1 or anti-PD-1, work by releasing the PD-1 receptor “brake” present on T cells. By preventing PD-1 from engaging PD-L1, a ligand expressed by tumor cells and tumor-infiltrating myeloid cells, these immunotherapeutic drugs suppress inhibitory signals transmitted to T cells. Currently nivolumab and pembrolizumab (anti-PD-1) and ipilimumab (anti-CTLA4) are approved by the US-FDA to treat metastatic MSI CRC.^5^

Comparatively, MSS CRCs which arise due to chromosomal instability display a comparatively weaker anti-tumor immune response, resulting in these tumors being largely refractory to this form of treatment.^6^ As MSS CRC accounts for the majority of advanced stage CRCs (~85%),^7^ sensitization of these tumors to immune checkpoint inhibition will represent a tremendous improvement in the therapeutic options available to these patients. Recent studies have identified a role for viral therapies as a promising, alternative strategy for cancer treatment. Respiratory enteric orphan virus (reovirus) is a double stranded RNA (dsRNA) virus consisting of a multilayered capsid protein structure. Our group and others have shown that reovirus preferentially replicates in *KRAS* mutant (*KRAS*^Mut^) cells,^8^ which account for approximately 45% of all CRCs,^9^ Reovirus enters *KRAS*^Mut^ cells through phagocytosis, un-coats capsid proteins in endosomes where it exploits the lower levels of eukaryotic translation initiation factor 2-α phosphorylation in these cells to enable viral dsRNA translation. This results in increased virion assembly, progeny generation and subsequent induction of apoptosis in infected cells.^10^

In addition to directly promoting oncolysis, oncolytic viruses (OVs) can also increase the recruitment of immune cells to an otherwise immunosuppressed TME to enhance antitumor effects. The antitumor immunity influenced by antiviral response is less clear across viral platforms and tumor types.^11^ After the entry and replication within tumor cells, the virus eventually lyses the cells and releases tumor antigens into the blood stream. These antigens can be detected by the immune system and draw T cells into the TME to initiate cancer cell killing, and potentially trigger the system to recognize metastatic disease elsewhere in the body.^12^ This opens up an avenue to combine reovirus treatment with ICIs in order to potentiate the efficacy of immunotherapy. Efficacy of both ICIs and OVs depends on factors such as cancer subtype, PD-1/PD-L1 expression, and the immune milieu.^13^

In metastatic CRC (mCRC), reovirus is currently in clinical development to treat *KRAS*^Mut^ as monotherapy or in combination with chemotherapy, however its potential to enhance the efficacy of immune checkpoint inhibitors has not been previously investigated.^8^ In this study, we tested the hypothesis that reovirus infection would lead to innate and adaptive immune responses and sensitize MSS CRC to PD-1 therapy, by administering reovirus as a single agent or in combination with an anti-PD-1 monoclonal antibodies in KRAS wild type (*KRAS*^Wt^) and mutant CRC cell lines, and in syngeneic mouse models of CRC.^14^ All cell lines were separated based on *KRAS* and mismatch repair (MMR) status (***Supplementary Table 1***).^15 16^ We observed significantly reduced tumor progression and increased survival of animals treated with the combination regimen and elucidate the mechanistic basis for this effect.

## Methods

### Cell culture and reagents

Fifty-nine human- and two mouse-origin CRC cell lines were obtained from range of sources.^15–17^ Human cell lines were cultured in MEM, CT26 in RPMI 1640, and MC38 in DMEM, all supplemented with 10% FBS, 10 mM L-glutamine, and penicillin (100 μg/mL)–streptomycin (0.1 mg/mL), at 37 °C under 5% CO2 pressure. Cell line authentication was performed using GenePrint^®^ 10 System and Fragment Analysis, and StemElite ID System (Promega, USA) at the Queensland Institute of Medical Research DNA Sequencing and Fragment Analysis Facility, Australia (January 2013) and the Genomic Core Facility, Albert Einstein College of Medicine, New York (June 2018).

### Reovirus infection

Reovirus type 3 dearing strain (Reolysin^®^) was provided by Oncolytics Biotech Inc. (Calgary, Canada). 0.5 to 2 × 10^6^ cells (depending on the experiment) were treated with reovirus multiplicity of infection (MOI) of 5. Cell were washed in PBS and harvested after 12 and 24 hours.

### Syngeneic *in vivo* models and allografts

Male and female BALB/c and C57BL/6 mice 6–8 week of age were purchased from Envigo Research Models & Services, NA Inc. All animal care and experimental procedures were performed in accordance with protocols approved by the Albert Einstein College of Medicine’s Institutional Animal Care and Use Committee (IACUC). BALB/c and C57BL/6 mice were intradermally injected with 10^5^ CT26 cells/mouse and 10^6^ MC38 cells/mouse, respectively, suspended in 100 uL PBS on the flank region. After the tumors had reached approximately 100 mm^3^ in size, mice were divided into four groups (n = 8-10/group) and treated with either reovirus intratumorally (i.t.) at a daily dose on 10 million tissue culture infective dose50 (TCID50) in 100 uL PBS [Reovirus group]; anti-mouse PD-1 (CD279) antibody (Clone: RMP1-14) intraperitoneally (i.p.) twice a week 200 ug in 100 uL PBS [anti-PD-1 group]; reovirus plus anti-PDI [Combination group]; or 100 uL PBS daily i.t. and isotype rat IgG2a κ 200 ug in 100 uL of PBS twice a week i.p. [Control group]. Tumor volume was measured every three days using calipers and calculated as follows: volume = longest tumor diameter × (shortest tumor diameter)^2^/2.^14^ Animals were euthanized and tumors were excised upon reaching a tumor volume of 2 cm^3^ size. For survival analyses, the health and behavior of the mice were assessed daily for the duration of the study. Upon presentation of defined criteria associated with tumor burden and disease progression (abnormal feeding behavior, diminished response to stimuli and failure to thrive), mice were humanely euthanized according to approved IACUC guidelines and survival time was recorded.

At the end of respective experiments, cultured cells and tumors were resected and either snap frozen or fixed in 10% buffered formalin for subsequent analyses.

### Cell viability assay

To determine reovirus sensitivity, 5,000-10,000 cells/well were seeded in 96-well plates. After 12-14 hours, cells were treated with reovirus at a 5 MOI for 24 hours. For each cell line, one plate was harvested at the time of viral infection for determination of t = 0 absorbance values. Viable cells were determined post treatment using the 3-(4,5-dimethylthiazol-2-yl)-2,5-diphenyltetrazolium bromide (MTT) (Sigma-Aldrich, USA; Cat# M2128) assay by measurement of absorbance at 570 nm.^17^ The relative rate of cell growth for each cell line was factored into the analysis by subtracting the absorbance at time 0 from both the control and treatment groups. All experiments were repeated at least three times, and each experiment was performed in technical triplicate

### Flow cytometry

To detect the expression of surface receptors and intracellular markers, cells were washed and incubated on ice for 20-40 minutes with appropriate fluorochrome-conjugated antibodies or isotype controls (***Supplementary Table 2***). Flow cytometric analysis was performed on a BD LSR II cell analyzer (BD Biosciences, USA), and the data analyzed using FlowJo version 9.1 (Tree Star, USA). For flow cytometric cell sorting, cells were stained with specific antibodies and separated on a BD FACSAria cell sorter (BD Biosciences, USA).^18 19^

### RT-qPCR

Total RNA was isolated from cell lines and tumor tissues using the RNeasy kit (Qiagen, USA) according to the manufacturer’s protocol. First-strand cDNA was synthesized using the iScript kit (Bio-Rad, USA). 10 ng of synthesized cDNA was used as template in real-time qPCR reactions using PowerUp SYBR Green (ThermoFisher, USA) on a Bio-Rad CFX96 RT-PCR machine. Changes in target gene expression were calculated using the 2-ΔΔCT data analysis method by comparing to the level of expression of GAPDH. Primer sequences are provided in ***Supplementary Table 2***.

### Transcriptome profiling using microarrays and computational analysis

Eighty nanograms of excellent quality RNA (RNA integrity number of ≥ 7.9) was hybridized on Clariom S gene expression arrays that interrogate the expression of > 20,000 transcripts. The study was carried out at the Genomic Core Facility, Albert Einstein College of Medicine. The arrays were scanned using a high-resolution GeneArray Scanner 3000 7G (ThermoFisher Scientific, USA). Data processing, normalization, background correction and array quality control were performed using Affymetrix’s Transcriptome Analysis Console (TAC). Annotation of probes was performed using the clariomshumantranscriptcluster.db package, and the average expression level was calculated for probes, which mapped to the same gene. For comparisons between groups, *Limma* package was used to perform an eBayes-moderated paired t-test provided in order to obtain log2 fold change (log2FC), p value, and adjusted p value (Benjamini-Hochberg-calculated FDR).^20^ Genes that displayed statistically significant tests (p value <0.05 and fold change [FC] ≥ 1.5/≤ −1.5) were considered differentially expressed (DEGs). Deconvolution analysis was performed using 847 immune-response related genes identified earlier by our group.^21 22^

Gene ontology (GO) terms and KEGG pathway enrichment analysis of DEGs: Analysis of GO terms such as biological process, cellular component, and molecular function were performed using the online tool DAVID (https://david.ncifcrf.gov) to identify systematic and comprehensive biological/cellular functions and conduct pathway exploration. The Functional Annotation setting in DAVID, which includes GO enrichment and use of the Kyoto Encyclopedia of Genes and Genomes (KEGG) pathways, helps to identify enriched genes among the list and uses previously annotated genes from the Clariom S array as background. GO categories with p value of <0.05 and FC ≥ 2/≤ −2 were considered to be significantly enriched.^23^

Protein–protein interaction (PPI) network analysis: To identify the hub regulatory genes and to examine the interactions between the DEGs, a PPI was generated using STRING (https://string-db.org/). These genes required an interaction score ≥ 0.2 and a maximum number of interactors = 0. The genes and corresponding PPI were imported into Cytoscape (version 3.6.1) with the Molecular Complex Detection (MCODE) app (version 1.5.1) to screen and visualize the modules of hub genes with a degree cut-off = 2, haircut on, node score cut-off = 0.2, k-core = 2, and max. depth = 100.^23^

### Transcriptome profiling using RNA-Seq

RNA-Seq analysis of 59 CRC cell lines was performed at the Australian Genome Research Facility (Melbourne, Australia) on an Illumina HiSeq2000 to a depth of >100 million paired reads as previously described by Mouradov et al., 2014.^24^ Absence of gene expression was defined as a RPKM value of <1.^25 26^

### Western blotting

Total cellular proteins were extracted using RIPA buffer and by solubilizing the proteins by boiling in SDS buffer (50 mM Tris-HCl, pH 7.5, 150 mM NaCl and 1% SDS). Protein lysates were then separated on an 8%-12% SDS-polyacrylamide gel. The proteins were then transferred to PVDF membrane (Millipore, USA) and incubated overnight at 4 °C with antibodies (***Supplementary Table 2***). Signal detection was performed using enhanced chemiluminescence (GE Healthcare, USA).^17^

### *Ex vivo* co-culture and real-time quantitative live-cell imaging

The IncuCyte S3 Live-Cell Imaging system (Essen Bioscience, USA^27^) was used for kinetic monitoring of cytotoxicity and apoptotic activity of CRC cell lines. CRC cells were seeded at a concentration of 4000 cells/well in a 96-well ImageLock™ plate (Essen BioScience), incubated overnight, and co-cultured with healthy human PBMCs at a ratio of 1:5 for 24 hours. Cells were transfected with Nuclight Green BacMam 3.0 Reagent and Cytotox Red Reagent (dead cell counting). After 24 hours, cells were treated with PBS, reovirus (2 MOI), anti-human PD-1 mAb (nivolumab; 2 nM), or a combination of reovirus and anti-human PD-1 (***Supplementary Table 3***). Plates were scanned and fluorescent and phase-contrast images were acquired in real-time in every 2 hours from 0 to 144 hours post treatment. Normalized green object count per well at each time-point and quantified time-lapse curves were generated by IncuCyte S3 2017A software (Essen BioScience). Cytotoxicity was calculated as Cytotox Red-positive cells divided by total cells per field, then divided by t = 0.^28^ The assays were performed on green-red overlay to quantify the number of cells that were dying through interactions of reovirus, anti-human PD-1, and their combination.

### Tumor microarray (TMA), immunohistochemistry (IHC) and immunofluorescence (IF)

Mouse tissue procurement: At the end of *in vivo* experiments, aliquots of tumor tissues were formalin fixed and paraffin embedded. Slides of tumor samples stained with hematoxylin and eosin were independently reviewed by a pathologist, and representative areas were marked. Core tissue biopsy specimens (2 mm in diameter) were obtained in triplicates from individual paraffin-embedded samples (donor blocks) and arranged in a new recipient paraffin block (tissue array block) using a tissue microarray construction punch needle (Newcomer Supply, USA). Each tissue array block contained up to 60 specimens, which allowed all 120 specimens (triplicate specimens of 20 cases) to be contained in 2 array blocks. Sections (4-μm) were cut from each tissue array block, placed on slides and deparaffinized, and dehydrated.

IHC was performed as previously described.^29^ In brief, antigen retrieval was performed by microwaving 4-μm sections in 0.01 M citrate buffer, pH 6.0, for 15 min at 650 W. Endogenous peroxidase activity was quenched with 3% hydrogen peroxide in methanol for 15 min. After incubating sections with blocking solution for 10 min, primary antibodies (***Supplementary Table 2***) were added at 4 °C for 12 h followed by biotinylated secondary antibody at room temperature for 10 min, and then streptavidin horseradish peroxidase (HRP) for 10 min. Staining was carried out with diaminobenzidine (DAB) chromogen, and counter-staining with Mayer’s hematoxylin. Blocking solution, secondary antibody, streptavidin HRP, and DAB were all purchased from the Cap-Plus Kit (Zymed Laboratories, USA). Stromal cells surrounding the tumor area served as internal positive controls.

Immunofluorescence was carried out using methods previously described.^30^ DAPI counterstain to identify nuclei and Cy-5-tyramide detection for target (C35, 1:500 dilution; Vaccinex, USA) for compartmentalized analysis of tissue sections were performed. Images of each TMA core were captured using a confocal microscope (Leica Microsystems, USA), and high-resolution digital images analyzed by the CaseViewer v.2.3 software (3D Histech, Hungary). While the number of tissues analyzed varied between different groups and species, all samples were in triplicate and n=7 per group as an average. We procured 2 sets of blocks per condition. Tumor staining was scored by a trained pathologist who had no knowledge about the data. The sections were scored semi quantitatively by light microscopy, using a 4-tier scoring system: 0, no staining; 1, weak staining; 2, moderate staining; 3, strong staining. In addition, the percentage of staining was also scored: 1 (0-25%); 2 (26-50%), 3 (51-75%) and 4 (76-100%). The final score for each section was obtained by multiplying the 2 scores.

### Statistical analysis

All data are presented as mean ± SEM of the indicated number of experiments. Differences were analyzed by Student’s t-test or ANOVA and Fisher’s post-hoc multiple comparisons test. Survival was analyzed with the Kaplan–Meier and log-rank tests for survival distribution. Results were considered significant when p < 0.05.

## Results

### Reovirus infection inhibits cell viability and induces key immune-response genes in CRC cell lines

To assess differential sensitivity of CRC cell lines to reovirus, thirteen CRC cell lines were infected with reovirus and cell viability assessed by MTT assay. The Caco-2 cell line (*KRAS*^Wt^, MSS) was found to be the most refractory to reovirus infection (75.5% cell survival after 24h) and HCT116 cells (*KRAS*^Mut^, MSI) the most sensitive (29.3% cell survival). Reovirus preferentially induced growth inhibition in *KRAS*^Mut^ cell lines (p=0.001), while no difference in sensitivity was observed between MSS and MSI lines (***Fig. 1A***).

**Fig. 1.**
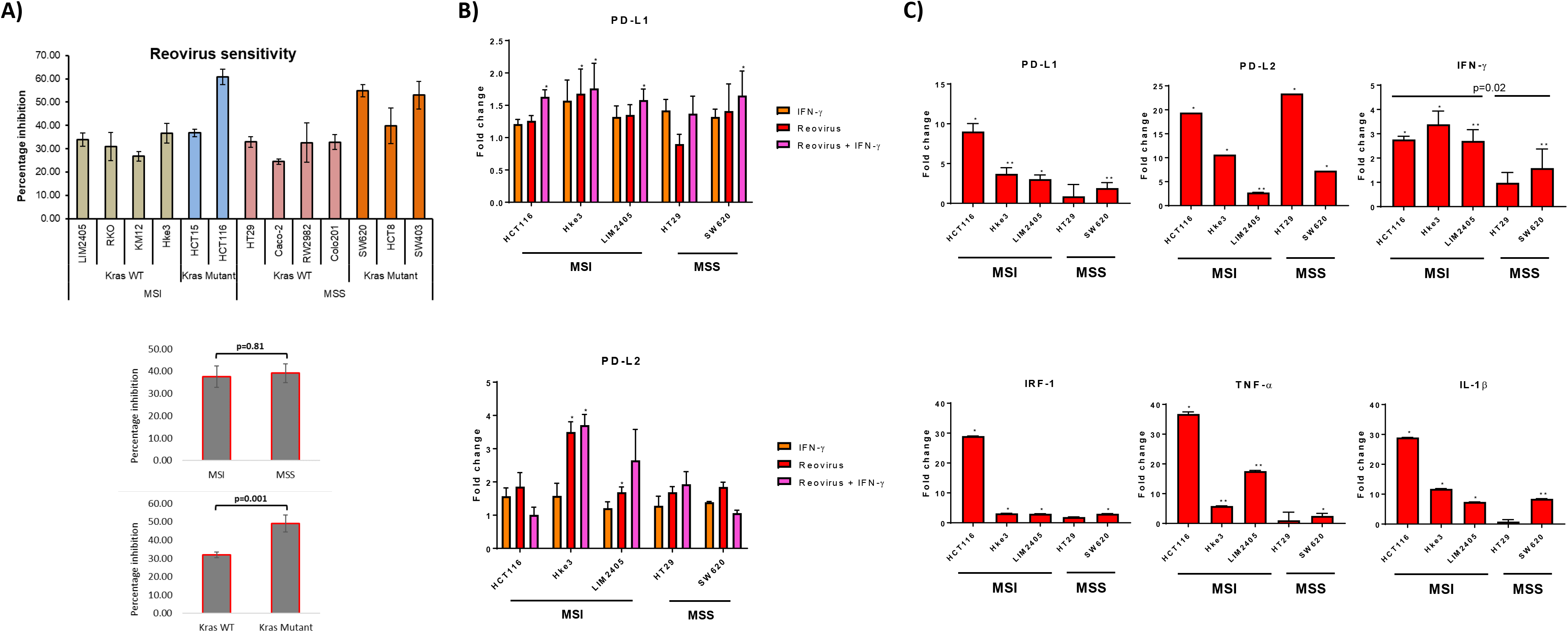
Measurement of cytotoxicity and expression of major immune response genes upon reovirus treatment. (A) MTT assay revealed that reovirus (reolysin) treatment of 5 MOI for 24 hours induced significant growth arrest in all 13 CRC cell lines studied. Mutational status of *KRAS* was more relevant than inherent MMR status of cell lines. (B) Flow cytometry analysis revealed that reovirus single agent and reovirus plus IFN-γ treatment increased PD-L1 expressing cell populations except for HT29. While reovirus treatment increased PD-L2 expressing populations among all cell lines, combination treatment with IFN-γ reduced *KRAS*^Mut^ cell lines such as HCT116 and SW620. (C) While transcriptional level expression of immune response regulators such as PD-L1, PD-L2, IFN-γ, IRF-1, TNF-α and IL-1β overall increased, no significant difference between *KRAS*^Mut^ and *KRAS*^Wt^ cell lines observed. MSI cell lines expressed significantly higher levels of IFN-γ compared to MSS cell lines upon reovirus infection. Most of the alterations were not prominent in HT29 expect for PD-L2.

We next assessed the effect of reovirus treatment on expression of immune-related genes. Cells were treated with reovirus (5 MOI) for 24 hours in the presence or absence of IFN-γ stimulation. Reovirus significantly increased PD-L1 and PD-L2 in presence of IFN-γ in all cell lines except HT29 (***Fig. 1B***). Similarly, RT-qPCR analysis showed significantly increased expression of immune response related genes, including PD-L1 and PD-L2, and also IFN-γ, IRF-1, TNF-α and IL-1β in presence of reovirus (***Fig. 1C***). HCT116 had 20 fold or more transcriptional level increase in PD-L2, IRF-1, TNF-α and IL-1β. Overall, 5 MOI reovirus treatment for 24 hours prompted significant growth reduction and expression of key immune-response genes among CRC cell lines.

### Bioinformatics analysis reveal inherent MSI/MSS status is more relevant than *KRAS* to regulate primary immune response to reovirus

To investigate the transcriptional reprogramming induced by reovirus infection, four CRC cell lines such as HCT116, Hke3, SW620 and Caco-2 were infected with reovirus and gene expression changes were assessed by microarray analysis 24 hour post-reovirus infection (***Fig. 2*** and ***Supplementary Fig. 1*)**. Hierarchical-clustering of reovirus-induced changes in expression of global immune-repose DEGs, and representation of pathways of viral immune response, TNF signaling and carcinogenesis under the classification of MMR (MSI/MSS) and *KRAS* mutational statuses were plotted. We observed increased number of DEGs under MSI/MSS classification when compared to *KRAS*^Mut^/*KRAS*^Wt^ (***Fig. 2A*** and ***Supplementary Fig. 1A***). The GO terms analysis revealed that TNF, NOD-like receptor (NLR), NF-κB signaling, and viral carcinogenesis pathways are associated with MMR status, and immune system process and osteoclast differentiation are with KRAS classification (***Fig. 2B*** and ***Supplementary Fig. 1B***). MSI, MSS, *KRAS*^Mut^ and *KRAS*^Wt^ PPI network from STRING database and hub genes of MMR and KRAS statuses network observed using the MCODE plugin in Cytoscape. These findings displayed the prominently upregulated genes that were involved in our samples. As well, to explore the functions of these genes, we looked for proteins that interact with the DEGs in STRING and constructed subsequent PPI networks (***Fig. 2C*** and ***Supplementary Fig. 1C***). The hub gene networks, along with the Venn diagram, also illustrated that cAMP responsive element binding protein 1 (CREB1) gene was highly upregulated under reovirus treatment in any circumstance or independent of genotypical characteristics (***Fig. 2D*** and ***Supplementary Fig. 1D***). Reovirus infection prominently activates programmed cell death among MSI, and innate immune response pathways among MSS cells, which were important regulators of NLR signaling,^31^ a function completely absent from the results of *KRAS*^Mut^ vs *KRAS*^Wt^ comparison under DAVID and STRING. In summary, specifically enriched GO terms such as positive regulation of immune system function, regulation of innate immune response, and innate immune response serve as indicators of involvement of T cell regulation^32^ under MSI/MSS classification. This supports the theory that immune-related genes are highly expressed within our samples and can be used towards the advantage of successful application of ICI as a partner drug to treat MSS type CRC.

**Fig. 2.**
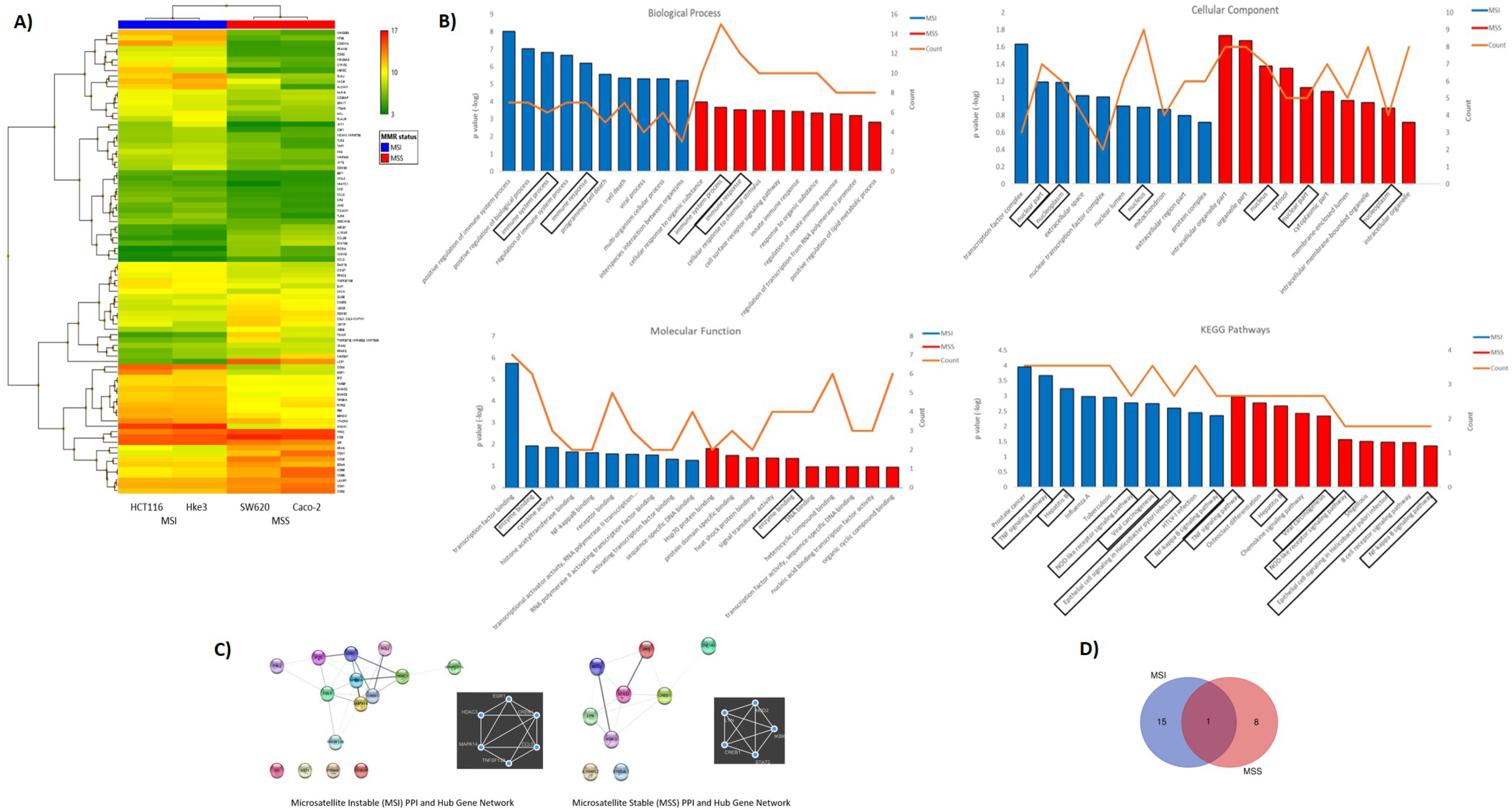
Differential expression of global immune response genes under reovirus infection in CRC cell lines categorized on MMR status. (A) Hierarchical-clustering profile across differentially expressed genes (DEGs) participating in immune pathways upon treatment with reovirus in MSI and MSS groups. RMA normalized expression values for the 86 genes were used to generate a heat-map in the TAC software. The colors indicate the expression value relative to the median expression value per gene in the dataset. Red indicates upregulation relative to median value and green indicates downregulation relative to the median value. (B) GO term enrichment and KEGG pathway analyses of DEGs using DAVID. Differentially expressed probes between MSI and MSS cell lines treated with reovirus were selected based on meeting the criteria of p ≤ 0.05, and divided into positive and negative fold changes lists and used to determine enrichment. (C) Protein interaction analysis using STRING. The different colored nodes represent different genes. Different edges (lines) show the interplay between genes. Edge strength is shown by thickness of the line, the thicker the line, the stronger the interaction. Hub networks made using MCODE demonstrate the genes within the PPI that were screened to display the most highly upregulated. The genes not included in the hub networks did not have substantial interaction amongst the other genes in the list, therefore they were not included. (D) Venn diagram created using the TAC software. It illustrates the clear point that between the MMR subgroups, CREB1 gene is the only commonality out of 25 genes, 16 in the MSI and 9 genes in the MSS groups.

### Selection of CRC cell lines based on stemness marker helps to establish *ex vivo* co-culture model

Next, we studied stemness markers in the previously analyzed cell lines, as these are associated with increased aggressiveness, tumor mass formation, and metastasis in CRC. In particular, we analyzed putative surface proteins CD133, CD44, CD24, and CD326/EpCAM (epithelial intracellular adhesion molecule)^33^ and connected the findings with their sensitivity to reovirus. Using RNA-Seq analysis, we studied the expression of stemness markers in 59 human CRC cell lines (***Fig. 3*** and ***Supplementary Fig. 2***). All 4 genes analyzed had well-defined expression in cell lines [RPKM >1 in at least 3/13 CRC cell lines by RNA-Seq] (total of n=20,702 genes). Nine cell lines that had abundant expression of stemness markers were chosen and validated using flow cytometry analysis (***Fig. 3A***). The cell lines LIM2405, HCT116, Caco2, HT29, SW620 and SW837 had increased expression of stem cell markers, with Caco2 that had low CD24 expression being an exception, and were selected for further studies. KM12, LIM1215 and RKO had overall reduced expressions of markers. In general, MSS had noticeably high expression of these markers compared to MSI.

**Fig. 3.**
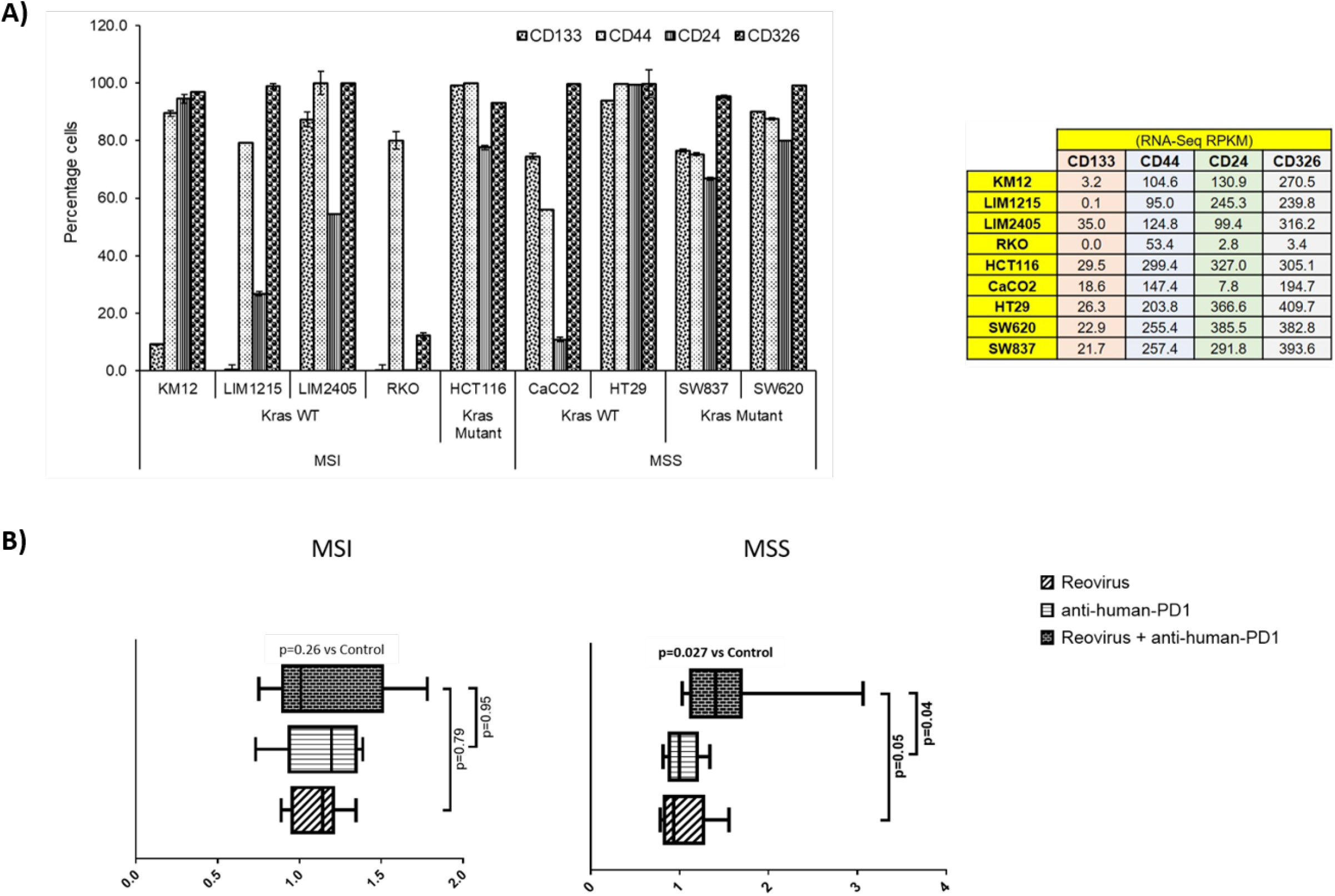
Selection of CRC cell lines, measurement of cytotoxicity, live-cell imaging and analysis in an *ex vivo* co-culture system. (A) Human CRC cell lines were selected based on expression of epithelial and stemness markers, and sensitivity towards reovirus infection. Out of 59 CRC cell lines screened using RNA-Seq (***Supplementary Fig. 2***), nine highly expressing ones (values are indicated on right panel table) with increased sensitivity to reovirus (***Fig. 1A***) were chosen and subjected to confirmatory analyses. One cell line per condition based on statuses of MMR and *KRAS* mutation with top expression of epithelial and stemness markers was taken for *ex vivo* co-culture studies (n=8). (B) Levels of cell death among MSI and MSS CRC cell lines co-cultured with human PBMC and treated with reovirus, anti-human PD-1 and their combination. Trends in fold difference and significance are depicted on box and whisker plot. Combination treatment compared to placebo rendered significant improvement in cell death only in MSS group. Combo increased cell death 1.6 fold in MSS and 1.2 fold in MSI (p values in bold letters indicate significance). Single agent treatments resulted in more than 1 fold cell death in all groups. Details of cell death was captured using live-cell imaging system and provided under ***supplementary movie*** section.

### Combinatorial administration of reovirus and anti-human PD-1 mAb increases cell death among MSS cell lines in the co-culture model

We next sought to determine whether reovirus treatment can enhance the anti-tumor efficacy of anti-PD-1. To do this, we performed *ex vivo* co-culture (1:5 CRC cells to healthy PBMC ratio) experiments using reovirus and anti-human PD-1 mAb and quantitatively determined cell death by 2 hourly live-imaging for 144 hours. MSS cells (SW620 and HT29) underwent significant cell killing in response to combinatorial therapy (2 nM nivolumab plus 2 MOI reovirus) when compared to control/placebo treatment (***Fig. 3B***). While single agent reovirus failed to show any significance compared to placebo or combination treatment, the difference between anti-human PD-1 administration and combination was significant. No significant difference was observed between anti-human PD-1 and placebo treatment among MSS cell lines. Further, only MSS cell lines collectively showed significantly increased cell death upon combination treatment (***Fig. 3B*** and ***Supplementary Fig. 3***). We performed phase-contrast live-cell imaging (Cytotox Red uptake) over a 144-hour period to characterize the kinetics of the cytotoxicity of reovirus and anti-human PD-1 towards CRC cell lines, HT29, SW620, LIM2405 and HCT116. The study demonstrated that combination-treated cells exhibit distinct cell morphology compared to untreated cells. Reovirus-treated cells showed membrane ballooning followed by membrane rupture while combination-treated cells displayed classical features of apoptosis such as cell shrinkage and membrane blebbing (apoptotic body formation). This unique morphology suggests combination-induced cell death is not distinct from apoptosis and is specific to HT29 (MSS) cells (***Supplementary Movie***).

### Differential effectiveness of reovirus and anti-PD-1 treatment in CT26 and MC38 syngeneic models of CRC

The effect of combining reovirus with an anti-PD-1 mAb on colon tumor growth was next examined *in vivo* using the CT26 (in BALB/c) and MC38 (in C57BL/6) syngeneic mouse models. In the *KRAS*^Mut^ CT26 model, single agent reovirus or anti-PD-1 treatment induced a modest decrease in tumor growth, however these effects were not statistically significant. Comparatively, combined treatment with reovirus and anti-PD-1 induced a highly significant suppression of the growth of established CT26 tumors. Consistent with the attenuation of tumor growth, combined reovirus/anti-PDL1 treatment significantly prolonged survival of mice harboring CT26 tumors. To assess the toxicity of reovirus and anti-PD-1 treatment, body weight was monitored throughout the treatment period and necropsy performed on one mouse from each group at the conclusion of the study (after 22 days of treatment) (***Fig. 4A***). Comparatively, no significant differences in tumor growth or survival were observed in the MC38 model.

**Fig. 4.**
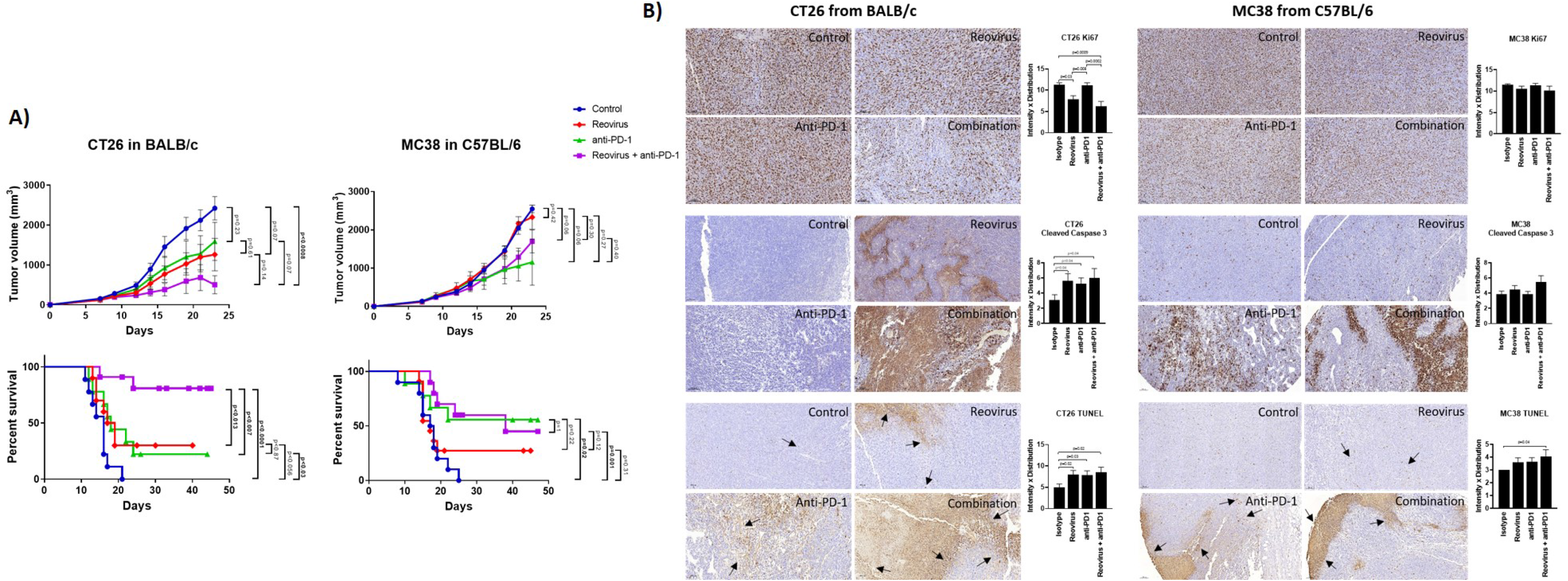
Effect of reovirus and anti-mouse PD-1 treatment on tumor growth, survival and expression of surface markers of proliferation and cell death in CT26 and MC38 syngeneic models. (A) In CT26, despite *KRAS*^Mut^ status, reovirus single agent couldn’t outperform the combination effects in terms of efficacy and survival. However, MC38 being MSI and strongly sensitive to ICI treatments, the combination therapy didn’t do any better than that of anti-PD-1 single agent. (B) Proliferation index, Ki67, and markers of apoptosis, cleaved caspase 3 and TUNEL were studied using IHC. While abundantly distributed and strongly stained, combination treatment profoundly reduced the expression of Ki67 in CT26 compared to MC38. Cleaved caspase 3 and TUNEL staining were more localized to apoptotic regions on tumors.

### Combinatorial treatment reduces the proliferative index and increases programmed cell death in CT26 tumors

To understand the mechanism driving the anti-tumor activity of the reovirus / anti-PDL1 treatment, resected tumors were stained for the proliferation marker, Ki67, and apoptosis markers cleaved-caspase 3 and TUNEL (***Fig. 4B***). Nuclear expression of Ki67 was decreased in reovirus and combination treated groups compared to isotype-treated controls in both models. However, the differences were significant only in CT26 model. Anti-PD-1 single-agent treatment didn’t affect the expression of Ki67 in either model. Expression of apoptosis indices significantly increased in the combination treated groups compared to those of controls. Reovirus single agent treatment selectively increased the expression of cleaved-caspase 3 and staining of TUNEL in CT26 compared to MC38. While anti-PD-1 administration alone increased TUNEL staining in both models, cleaved-caspase 3 expression significantly increased only in CT26. Although increased evidence was noticed in CT26, combination treatment had profoundly increased the expression of these markers in both models. Ki67 staining was distributed unevenly across all samples and conditions. The distribution of apoptotic markers were more specific to tumor-rich areas in CT26, whereas it was wide-spread and unevenly dispersed in MC38 under the combination treatment.

### Mixed alterations of PD-L1/PD-1 axis by reovirus and anti-PD-1 treatment on CT26 and MC38 tumors

To examine whether treatment with anti-PD-1 alone or in combination with reovirus impacted the PD-1/PD-L1 axis, we examined expression of components of this pathway in the resected tumors. Expression of PD-L1, the key ligand of PD-1, was significantly increased upon reovirus and reovirus plus anti-PD-1 administration (p=0.03; n=5) in CT26 tumors. MC38 tumors had reduced expression of PD-L1 under all treatment condition. While anti-PD-1 administration reduced the expression of PD-1 protein in both CT26 and MC38 tumors, only CT26 had further reduction upon combining with reovirus (p=0.01; n=5). Indeed, reovirus single agent and combination groups had increased expression of PD-1 in MC38 (***Fig. 5A*** and ***5B***, & ***Supplementary Fig. 5A*** and ***5B***).

**Fig. 5.**
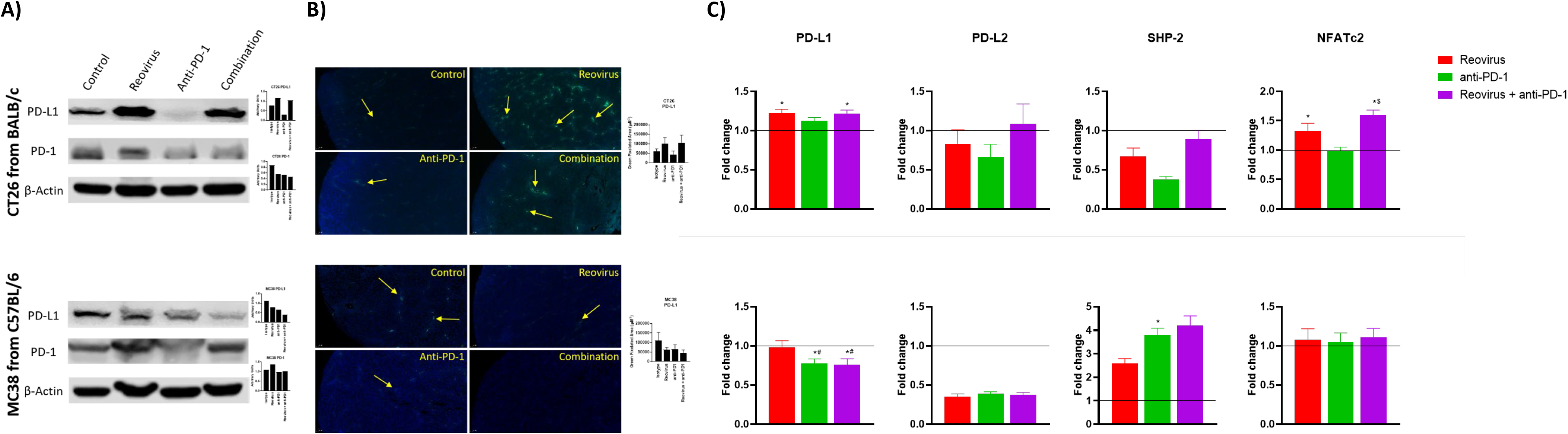
Differential activation of T cells via PD-L1/PD-1 signaling in CT26 and MC38 tumors. (A) Single agent reovirus treatment increased (p=0.04) or unaltered of PD-L1 expression and unaltered or increased (p=NS) of PD-1 in CT26 and MC38 tumors, respectively. Trends in expression also reversed in both models upon combination treatment, however, significance observed only for PD-L1 (p=0.03; CT26, and p=0.04; MC38). (B) Representative immunofluorescence images of PD-L1 expression among CT26 and MC38 tumors. PD-L1 expression significantly increased upon reovirus (p=0.03) and its combination with anti-PD-1 treatment (p=0.04) in CT26. While the expression was unaltered with single agent treatments, combination reduced (p=0.05) PD-L1 in MC38 tumors. (C) Transcriptional level changes in expression of key mediators of PD-L1/PD-1 signaling. Combination treatment increased the fold difference in expression of PD-L1 (p=0.04), PD-L2, SHP-2 and NFATc2 (p=0.03) in CT26 and SHP-2 in MC38. Combo decreased PD-L1 (p=0.03) and unaltered PD-L2 and NFATc2 expression in MC38 tumors. Fold difference was comparatively low for all genes investigated under single agent anti-PD-1 treatment in CT26, and PD-L1 alone in MC38, where the expression indeed reduced (p=0.03). Data were shown as mean ± SEM; n = 5; * p ≤ 0.05, compared with control group; # p < 0.05 compared with reovirus group; $ p ≤ 0.05, compared with anti-PD-1 group.

SHP-2, the protein tyrosine phosphatase that mediates the negative costimulatory role of PD-1, was also reduced by the absence of PD-1 in CT26 tumors under reovirus, anti-PD-1 and combination administration. SHP-2 levels went up by all treatments, particularly, by anti-PD-1 treatment in MC38. Interestingly, NFATc2, one of the transcription factors activated by T cell receptor stimulation increased upon treatment with reovirus and its combination with PD-1 in CT26 tumors. There was no change in expression of NFATc2 observed under any treatment conditions in MC38 tumors (**Fig. 5C**). In summary these studies reveal differential activation of PD-L1/PD-1 axis in the TME of CT26 and MC38 tumors upon treatment with reovirus and anti-PD-1.

### Reovirus and anti-PD-1 combination therapy synergistically enhances the anti-tumor adaptive immune response

Next, we analyzed the composition and activation status of tumor-infiltrating immune cells in the CT26 and MC38 models treated with this combination. We observed increased T cell infiltration in both MC38 and CT26 tumors treated with anti-PD-1 or reovirus alone, with a further increase in tumors treated with the combination (***Fig. 6A***, top). Similarly, TILs were increased in CT26 tumors treated with anti-PD-1 or reovirus alone and further increased by the combination therapy. In contrast, TILs weren’t significant among MC38 tumors under any treatment modality (***Fig. 6A***, bottom). Lastly, the combination therapy increased granzyme B expression more than either monotherapy in CT26 tumors, whereas this synergistic response was not seen in MC38 tumors (***Fig. 6B***). This indicates that anti-PD-1 and reovirus combination therapy enhances a cytotoxic immune response in the MSS CT26 model to a greater degree than either monotherapy.

**Fig. 6.**
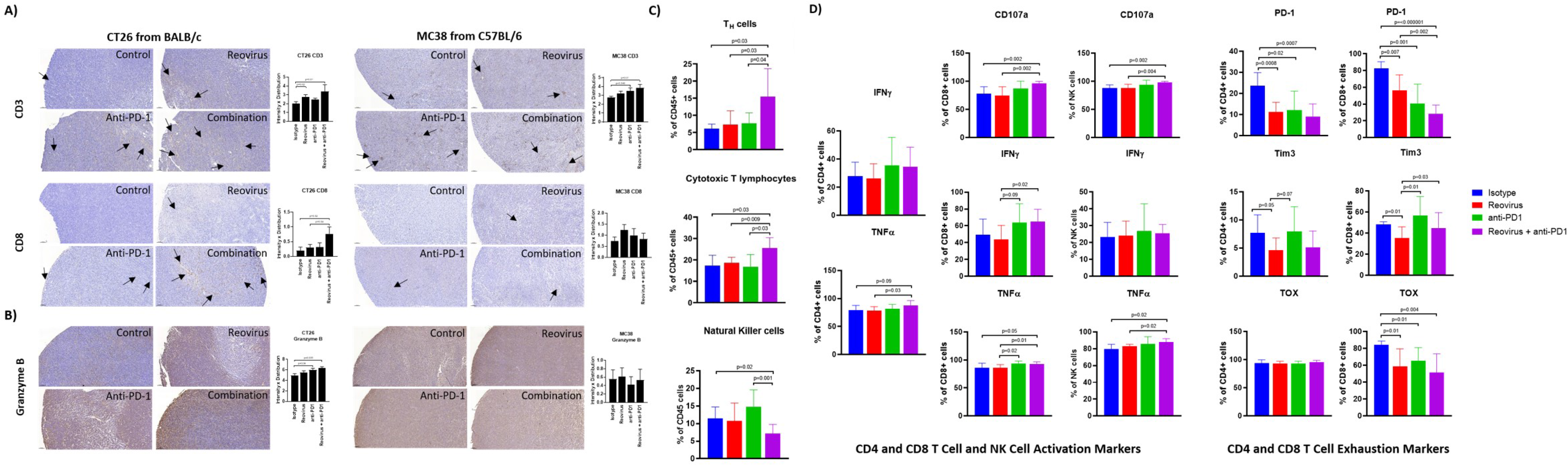
Difference in localization, distribution and expression of cell surface and activation/exhaustion markers in CD4^+^ and CD8^+^ T cell populations upon reovirus and anti-PD-1 treatment. (A) Reovirus in CT26, anti-PD1 in MC38, the combination treatment in both models increased the expression of CD3 (p=NS vs control). While overall, weak distribution, CD8^+^ T cell staining was increased upon reovirus (p=0.04) and combination treatment (p=0.03) specifically in CT26. (B) Granzyme B, which is a functional marker of NK cells also, increased upon treating with reovirus (p=NS) and combination (p=0.03) in CT26. Overall granzyme B distribution, while high, we didn’t observe any significant difference among groups in MC38. (C) While single agent treatments displayed insignificant role, combo increased CD4^+^ and CD8^+^, and reduced NK cells among overall CD45^+^ cell populations in CT26. Strikingly enough, the increment among NK cells boosted by anti-PD-1 abrogated by reovirus administration (p=NS with reovirus). (D) Despite overall activation of CD4^+^ and CD8^+^ T cells, significantly increased expression of surface markers (CD107a, IFN-γ and TNF-α) observed only among CD8^+^ cells upon treatment with reovirus plus anti-PD-1. Anti-PD-1 single agent treatment seemingly increasing the expression of all markers, except TNF-α in CD4^+^ T cells. While anti-PD-1 alone increased CD107a, IFN-γ and TNF-α expression (p=NS) among NK cells, combination treatment significantly increased only CD107a. Reovirus treatment didn’t make differences in expression in any of the cytokines studied among NK cell populations. (E) Single agents and combination treatment largely reduced the expression of exhaustion markers, however, CD4^+^ TOX among all groups and Tim3 among CD4+ and CD8^+^ T cells after anti-PD-1 treatment. Noticeably, Tim3 expression among CD8^+^ T cells were increased upon anti-PD-1 treatment.

To obtain a more detailed and comprehensive view of the TME in the CT26 model, we enzymatically dissociated the tumors and performed flow cytometric analysis. Consistent with our findings obtained by IHC staining (***Fig. 6A***), we found that the combination treatment synergistically enhanced the infiltration of both CD4^+^ and CD8^+^ T cells (***Fig. 6C***). We did note a relative decrease in the proportion of CD49b^+^ NK cells (***Fig. 6C***), although this is likely due to the increased proportions of T cells, DCs, and monocytes in these tumor microenvironments (***Fig. 7B***). We next analyzed the functional state of the tumor-infiltrating T cells and NK cells. We found that the combination treatment increased the intracellular expression of the inflammatory cytokines IFN-γ and TNF-α, as well as the cytotoxicity marker CD107a in CD8^+^ T cells, whereas anti-PD-1 or reovirus monotherapy did not notably increased these markers (***Fig. 6D***). Likewise, the combination therapy also increased the expression of TNF-α in CD4^+^ T cells as compared to either monotherapy, although it did not have a marked effect on IFN-γ in these cells. The combination treatment also enhanced cytotoxicity and TNF-α expression in NK cells (***Fig. 6D***). The increased expression of TNF-α in CD4+ T cells was also compounded with a significant decrease in the expression of inhibitory cytokine TGF-β in the combinatorial treatment of CT26 tumor cells when compared to single agents further confirming the immune stimulation (***Supplementary Fig. 4***).

**Fig. 7.**
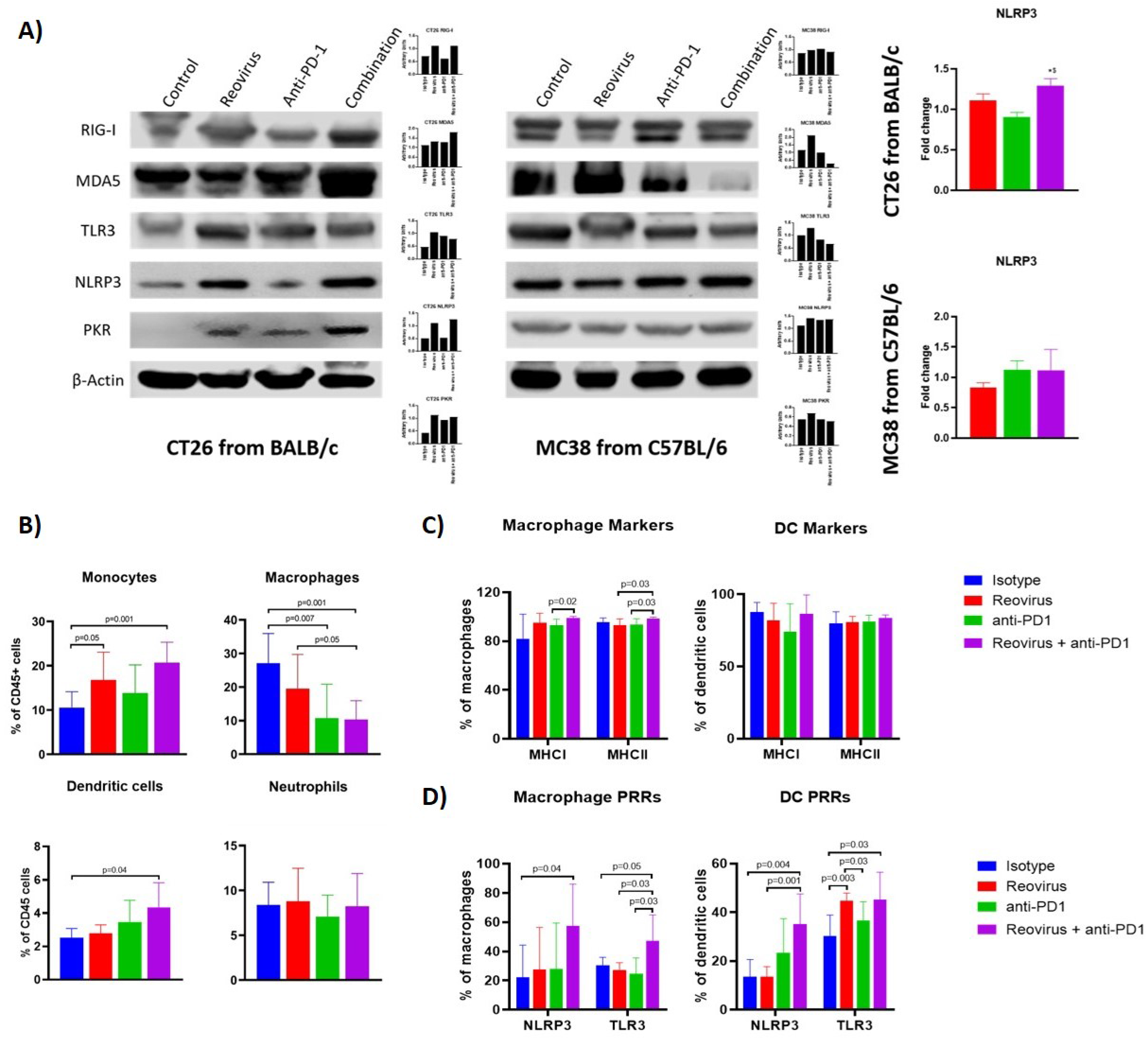
Activation of pattern recognition receptors (PRRs) and innate immune system in CT26 and MC38 tumors treated with reovirus, anti-PD-1 and their combination. (A) Protein-level changes among pooled samples - reovirus infection upregulated most of the PRRs, except for cytoplasmic RIG-I, in MC38. Combination treatment increased almost all PRRs, except TLR3 and PKR, in CT26, and either unaltered or decreased in MC38. Combo drastically reduced membrane bound TLR3 and cytoplasmic PKR in MC38. NLRP3, which is an inflammasome activation mediator, significantly increased by the effect of reovirus in the TME of CT26. Expression of dsRNA-sensing PRR, PKR, was increased by reovirus treatment in both models combo, however, didn’t change in expression in MC38. (B) Effect of reovirus and anti-PD1 treatment on innate immune cell populations. Reovirus single agent and combo treatment increased monocytes and DCs (p=NS for DCs with reovirus), and decreased macrophages. Anti-mouse PD-1 treatment alone increased percentages of monocytes and DCs in the TME. (C) Differential expression of antigen presentation markers on the surface of macrophages and dendritic cells. Reovirus or combo treatment didn’t alter MHC Class I and II expression significantly among these cell populations. Only macrophage MHC I and II expression significantly increased in combo compared to anti-mouse PD-1 single agent. (D) Selective activation of PRRs in tumor-derived myeloid cells. While combination treatment did in all groups, neither of the single agents significantly altered PRRs expression, except anti-PD-1 and reovirus on TLR3 among DCs. NLRP3 and TLR3 expression almost tripled among cell types by the combination treatment.

As CD8^+^ T cells encounter persistent stimulation and immunosuppressive signals in the TME, they can acquire a dysfunctional or “exhausted” state.^34^ Anti-PD-1 therapy has been reported to relieve T cell exhaustion,^35^ therefore we hypothesized that combination therapy with reovirus would synergistically decrease T cell exhaustion in infiltrating TILs. To analyze T cell exhaustion, we analyzed the proportion of T cells that expressed the exhaustion markers PD-1 and Tim-3, as well as the transcription factor TOX that has been recently reported to be a specific marker for terminal exhaustion in CD8^+^ T cells. Intriguingly, we found that the combination therapy potently reduced T cell exhaustion in CD8^+^ T cells, as seen by reductions in both PD-1 and TOX expression (***Fig. 6E***). Although significance was not reached, the regulatory T cell (Treg) population was reduced in the combination group as compared to the anti-PD1 further indicating immune activation (***Supplementary Fig. 4***). Collectively, our data show that reovirus and anti-PD-1 combination therapy synergistically promotes CD8^+^ and CD4^+^ T cell infiltration, enhances inflammatory and cytotoxic effector functions, and reduces CD8^+^ T cell exhaustion.

### Selective activation of pattern recognition receptors (PRRs) and expression of antigen presentation markers of innate immune cell types by reovirus and anti-PD-1 combination

Activation of innate immune cells, particularly antigen-presenting cells such as dendritic cells and macrophages, are key to an effective anti-tumor immune response. We hypothesized that reovirus infection would promote the activation of innate immune pathways through PRR pathways. Sensing of the dsRNA that is produced as a replicative intermediate during reovirus infection is done by host PRRs such as retinoic acid-inducible gene I (RIG-I), melanoma differentiation-associated protein 5 (MDA-5), toll-like receptor 3 (TLR3) and NOD-like receptor protein 3 (NLRP3) leading to the induction of the innate immune response via the type I interferon pathway.^36^ The effects of reovirus or anti-PD-1 therapy on PRRs was studied in CT26 and MC38 tumors by Western blotting (individual and pooled samples) and qPCR (***Fig. 7A; Supplementary Fig. 5A*** and ***5B***). Cytoplasmic PRRs such as RIG-I and MDA5 and endoplasmic or plasma membrane-bound TLR3 were increased in CT26, and unaltered or decreased in MC38 - particularly, MDA5 - after treatment with reovirus and anti-PD-1. TLR3, NLRP3 (inflammasome mediator PRR), and protein kinase R (PKR; cytoplasmic sensor of viral-mediated dsRNA) were upregulated under both single agent reovirus- and combination-treated groups in CT26, however, the changes were insignificant in MC38 (p > 0.05). Notably, the increased expression of RIG-I, MDA5, PKR, and NLRP3 was greatest in the combination treatment group, suggesting a synergistic increase in the activation of innate immune pathways. We further confirmed the increased expression of NLRP3 at the mRNA level with qPCR (***Fig 7A***, right).

To further analyze the composition of the tumor-infiltrating myeloid cells, we performed flow cytometric analysis of dissociated tumors. We found that the combination treatment significantly decreased the populations of CD11b^+^ F4/80^+^ macrophages, which are a major population of immunosuppressive cells in the TME. In contrast, the combination treatment increased the populations of Ly6C^+^ monocytes and CD11c^+^ dendritic cells (DCs). No significant changes were seen in the Ly6G^+^ neutrophil population in any treatment group (***Fig. 7B***). Next, we sought to analyze the expression of activation markers and PRRs in the myeloid cells. The expression of antigen-presentation molecules MHCI and MHCII on macrophages and DCs was high at baseline^19^ even in the isotype-treated group and was not significantly increased with treatment (***Fig. 7C***). Interestingly, expression of the PRRs such as NLRP3 and TLR3 was increased with combination treatment in both macrophages and DCs (***Fig. 7D***). This indicates that the increased expression of PRRs seen with Western blotting was at least in part due to the contribution of tumor-infiltrating myeloid cells. Put together, our studies show that combination reovirus and anti-PD-1 treatment synergistically activates innate immune PRR pathways, reduces immunosuppressive macrophage populations while promoting effector monocyte and DC populations, and increases the expression of PRR molecules within both macrophages and DCs.

## Discussion

The KEYNOTE-016 trial that evaluated pembrolizumab monotherapy, and CheckMate-142 trial which evaluated nivolumab plus ipilimumab examined response of MSI mCRC to immunotherapy, but also included a subset of patients with MSS mCRC.^37 38^ In contrast to the positive results observed in MSI tumors, MSS mCRC patients did not respond to checkpoint inhibition, highlighting the predictive value of microsatellite instability in response to immunotherapy in mCRC. Thus, it is of critical importance to find strategies to improve the response of immune cold tumors to ICI. Here, we demonstrate the feasibility and efficacy of using oncolytic reovirus therapy alongside anti-PD-1 treatment as a novel combination treatment strategy for MSS CRC tumors.

Using *in vitro* experiments, microarray, and bioinformatics analysis of global immune-response genes, we highlighted a first-of-its-kind discovery that the inherent MMR status (MSI or MSS) is more powerful influencer than *KRAS* mutational status to determine sensitivity to reovirus infection. This prompted us to formulate the hypothesis that a combination of reovirus with a clinically approved ICI, anti-PD-1, would have synergistic therapeutic efficacy against MSS type of CRC tumors. Heavily pretreated CRC patients were administered with the oncolytic vaccinia virus (Pexa-Vec [JX-594] engineered to express GM-CSF, a hematopoietic growth factor that increases dendritic cell differentiation, maturation and function and induced tumor reactive T cells^39^ and β-galactosidase) and reached stable disease in 67% (n = 10) of patients. The biweekly injection did not lead to dose-limiting toxicities in this phase Ib study alone^40^ or in a phase I/II study in combination with checkpoint inhibitors tremelimumab (CTLA–4) and durvalumab (PD-L1).^41^ A phase Ib trial testing the combination of T-VEC and pembrolizumab revealed a high overall response rate (ORR) of 62% and complete responses in 33% of melanoma patients independent of baseline CD8^+^ infiltration.^42^ These mixed results reveal the hurdles that oncolytic viral therapy must still overcome. Some of these challenges include optimizing tumor tropism, viral delivery, and enhancing anti-tumor immunity.

We demonstrate that the antitumor activity of the drug combination is associated with both the induction of apoptosis in tumor cells as well immune activation. With regards to the former, our time-lapse live-cell imaging and IHC data indicate that combinatorial treatment reduced the expression of Ki67 while enhancing the expression of cleaved-caspase3 and TUNEL staining. Our group and others have demonstrated that defective activation of the antiviral PKR-eIF2α pathway is a key determinant of direct oncolysis initiated by reovirus infection,^10 17 43 44^ enabling efficient viral replication, and immune activation in a later stage.^22^ Reovirus infection also triggers the cellular interferon (IFN) response to produce Type 1 IFN’s alpha and beta (IFNα/β). Secreted IFNα/β can stimulate the JAK-STAT pathway in an autocrine or paracrine manner to activate hundreds of IFN-stimulated genes (ISGs), many of which have antiviral activities that elicit an antiviral state.^45^

Activation of immune response KEGG pathways such as, TNF-, NOD-like receptor-, or NF-kB-signaling following reovirus treatment could then further enhance the activity of this combination by enhancing immune-mediated cell killing, as observed in in the co-culture system.^46–48^ In addition to these pathways, additional pathways may also be involved. For example, we observed that reovirus treatment increased expression of CREB1 in both MSI/MSS and *KRAS*^Mut^/*KRAS*^Wt^ cell lines under reovirus treatment indicates altered cell death signaling and viral immune mediation.^49^ CREB1 has been shown to play a large role in TNF signaling pathways,^50^ raising the possibility reovirus treatment may increase TNF signaling within CRC tumors, leading to apoptosis and total cell death.^51^ Combination of agents that increase the expression of death receptor 5 (DR5) and its ligand, TNF-related apoptosis-inducing ligand (TRAIL), is a novel anti-CRC therapy, and correlate with tumor regression.^52^ Alongside reovirus treatment, TNF signaling upregulation secondary to CREB1 expression may prove efficacious to CRC treatment in future studies. Targeting CREB1 binding protein/β-catenin, combined with PD-1/PD-L1 blockade, has shown potential as a new therapeutic strategy for treating liver metastasis in CRC.^53^ Further studies are needed on the role of CREB1 and TNF signaling in MSS cancers and their susceptibility to immunotherapy.

Consideration of treatment regimens will be particularly important for combination of reovirus with ICIs because anti-CTLA-4 antibodies are likely to potentiate early stages of T cell priming, whilst anti-PD1/anti-PD-L1 mAbs would act to reverse T cell exhaustion within the TME.^54^ In a recent combinatorial study, reovirus-specific – but, not tumor-specific – CD8^+^ TILs served as non-exhausted effector cells for the subsequently systemically administered CD3-bispecific antibodies.^55^ OV-infected cancer cells tend to down-regulate their class I MHC making themselves a good target for functionally active CD107a^+^ NK cells. Although NK cells may kill infected cancer cells and limit the amplification of OVs, studies have found that NK cells often have positive effects on therapeutic outcomes of OVs.^56 57^ Proving the oncolytic nature of reovirus and its influence on immune response mediators such as cytokines and cellular regulators are important in determining its possibility to combine with an ICI to test synergistic efficacy. The activation of innate-sensing pathways of antigen-presenting CD11c^+^ DCs, CD11b^+^ macrophages and monocytes within TME may trigger enhanced CD8^+^ T cell responses against the tumor.^58^

OVs promote PD-L1 expression in both cancer and TME cells and by effectively combining with an anti–PD-L1 antibody this barrier can be effectively overcome.^59^ Increased effect of reovirus by anti-PD-1 treatment was analyzed using double-stranded RNA-dependent kinase - PKR - that induces inflammation by regulating the expression of the NLR family pyrin domain-containing 3 inflammasome – NLRP3 - through NF-κB signaling. NLRP3 and TLR3 sense a wide range of pathogen-associated molecular patterns (PAMPs) and damage-associated molecular patterns (DAMPs).^60^ We observed an increase in the expression of NLRP3 in CT26 CRC TME which corroborates and augments the previously published findings regarding the function of NLRP3. APCs expressing PRRs such as TLR3, MDA5, RIG-I, NLRP3 etc. can be directly activated by PAMPs or DAMPs to become competent to prime T cell responses.^36^ Engagement of PRRs on DCs induces NF-κB activation, up-regulation of co-stimulatory molecules, and production of cytokines and promotion of cross-priming.^61^ PAMPs and DAMPs can also be produced by immunogenic cell death (ICD) of tumor cells induced by reovirus. We are thus confident that reovirus and anti-PD-1 antibody combination is a promising therapeutic approach to convert cold MSS tumors to inflamed immune therapy sensitive tumors. Thus, reovirus promotes an overall inflammatory TME and increases the responsiveness of MSS tumors to immunotherapy.^62^

## Conclusion

Herein we report for the first time that immune insensitive MSS tumors that account for 85% of all CRC can be effectively converted to immunotherapy sensitive cancers by introducing reovirus. This has successfully immune populated the MSS tumor TME in our three study models. Immunotherapy that has achieved phenomenal success in many cancer types may now be successfully implemented in MSS ‘cold’ or immunosuppressive CRC tumors, and provides a rationale to extend combination reovirus-ICI therapy into clinical testing.

## Declarations

## General and Funding

S. Goel was supported by NIH/AG1R21 NIH grant AG 1R21AG058027-01. R. Maitra was supported by Yeshiva University startup fund 2A4108. We gratefully acknowledge the Office of Grant Support, Genomic Facility, Flow Cytometry Core Facility, and the Analytical Imaging Facility of Albert Einstein College of Medicine along with the NCI cancer center support grant (P30CA013330), which partially supports all morphometric work conducted with 3D Histech P250 High Capacity Slide Scanner SIG #1S10OD019961-01 of the shared facilities. We also thankfully acknowledge Dr. Lidija Klampfer of Southern Research Institute, Birmingham, Alabama for the kind gift of Hke3 cell line.

## Author Contributions

**T. Augustine**: Conceptualization, supervision, resources, methodology, investigation, analysis, and writing the original draft. **P. John**: Methodology, investigation, formal analysis, validation, data curation, and writing in-part. **T. Friedman**: Methodology, investigation, formal analysis, methodology, software, and writing in-part. **J. Jiffry**: Methodology, investigation and formal analysis. **H. Guzik**: Methodology, investigation, and visualization. **R. Mannan**: Formal analysis, and investigation. **R. Gupta**: Formal analysis, and investigation. **C. Delano**: Methodology, formal analysis, and investigation. **J.M. Mariadason**: Supervision, methodology, analysis and editing. **X. Zang**: Supervision, methodology, analysis and editing. **R. Maitra**: Data curation, supervision, methodology, analysis, and editing. **S. Goel**: Conceptualization, resources, data curation, software, formal analysis, supervision, funding acquisition, validation, methodology, project administration, and editing.

## Supplementary Materials

**Supplementary Table 1.** Molecular profiling of key CRC cell lines studied in the manuscript. Summary of microsatellite instability and *KRAS* gene mutation statuses of seven human and two mouse CRC cell lines used in the manuscript.

|  | Cell Line | MMR status | *KRAS/Kras* status |
| --- | --- | --- | --- |
| Human | HCT116 | MSI | MUT |
|  | Hke3 | MSI | WT |
|  | LIM2405 | MSI | WT |
|  | HT29 | MSS | WT |
|  | SW837 | MSS | MUT |
|  | SW620 | MSS | MUT |
|  | Caco2 | MSS | WT |
| Mouse | MC38 | MSI | WT |
|  | CT26 | MSS | MUT |

\*MSI – microsatellite unstable, MSS – microsatellite stable, MUT – mutant, WT – wild type

**Supplementary Table 2.**
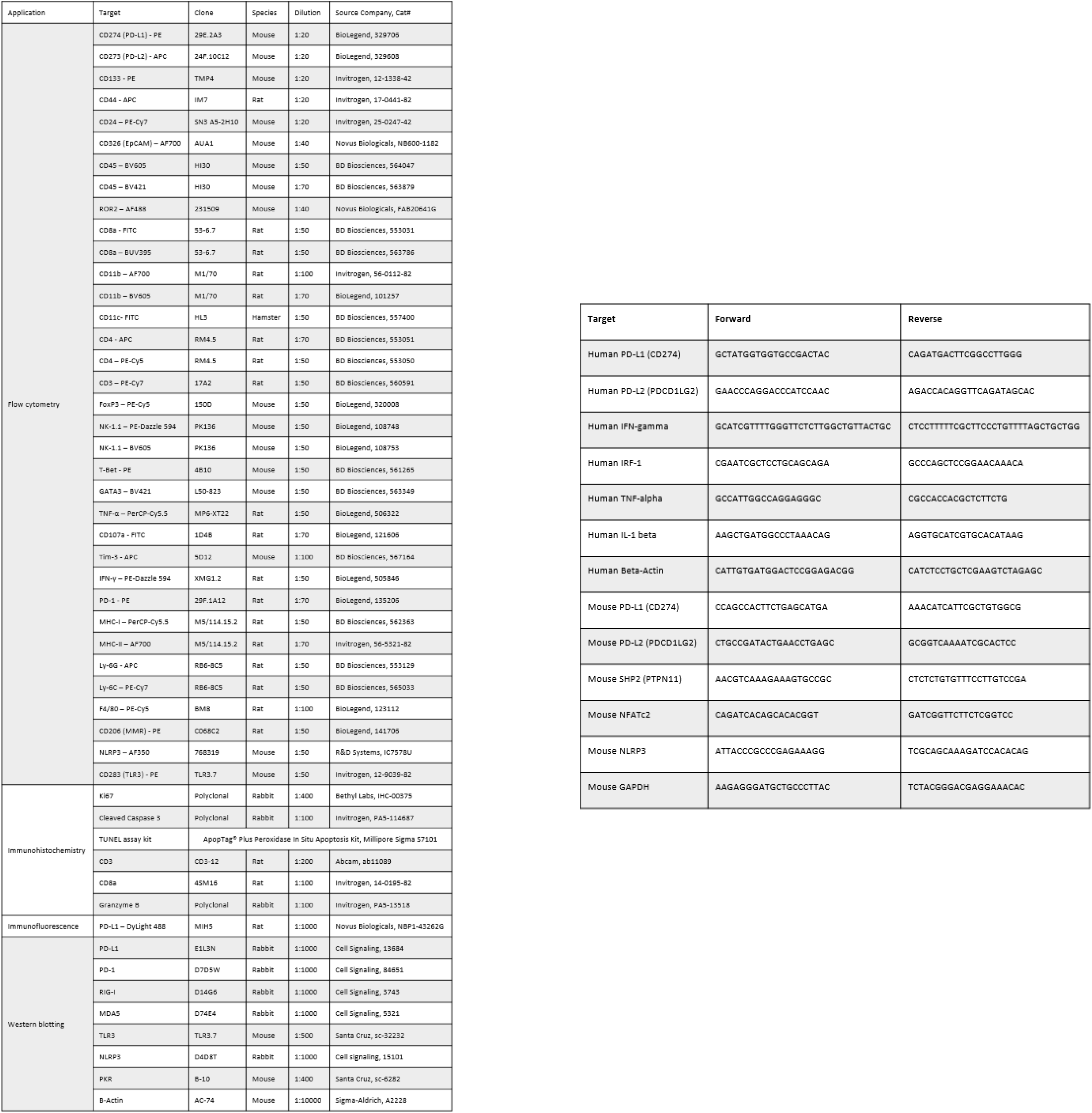
List of antibodies and primers used.

**Supplementary Table 3.** Details of the experimental set-up of *ex vivo* co-culture study. Every group was added with human CRC cell lines and human PBMCs consecutively as part of establishing the culture, and nuclear labeling and dead cell counting reagents for identification and measurement, respectively.

| Group | Experimental Set-up | Group |
| --- | --- | --- |
| 1 | CRC cells + 5X PBMC + BacMam Green + Cytotox Red + PBS | Control |
| 2 | CRC cells + 5X PBMC + BacMam Green + Cytotox Red + 2 MOI Reovirus | Reovirus |
| 3 | CRC cells + 5X PBMC + BacMam Green + Cytotox Red + 2 nm Nivolumab | Anti-human-PD-1 |
| 4 | CRC cells + 5X PBMC + BacMam Green + Cytotox Red + 2 MOI Reovirus + 2 nm Nivolumab | Combination |

**Supplementary Fig. 1.**
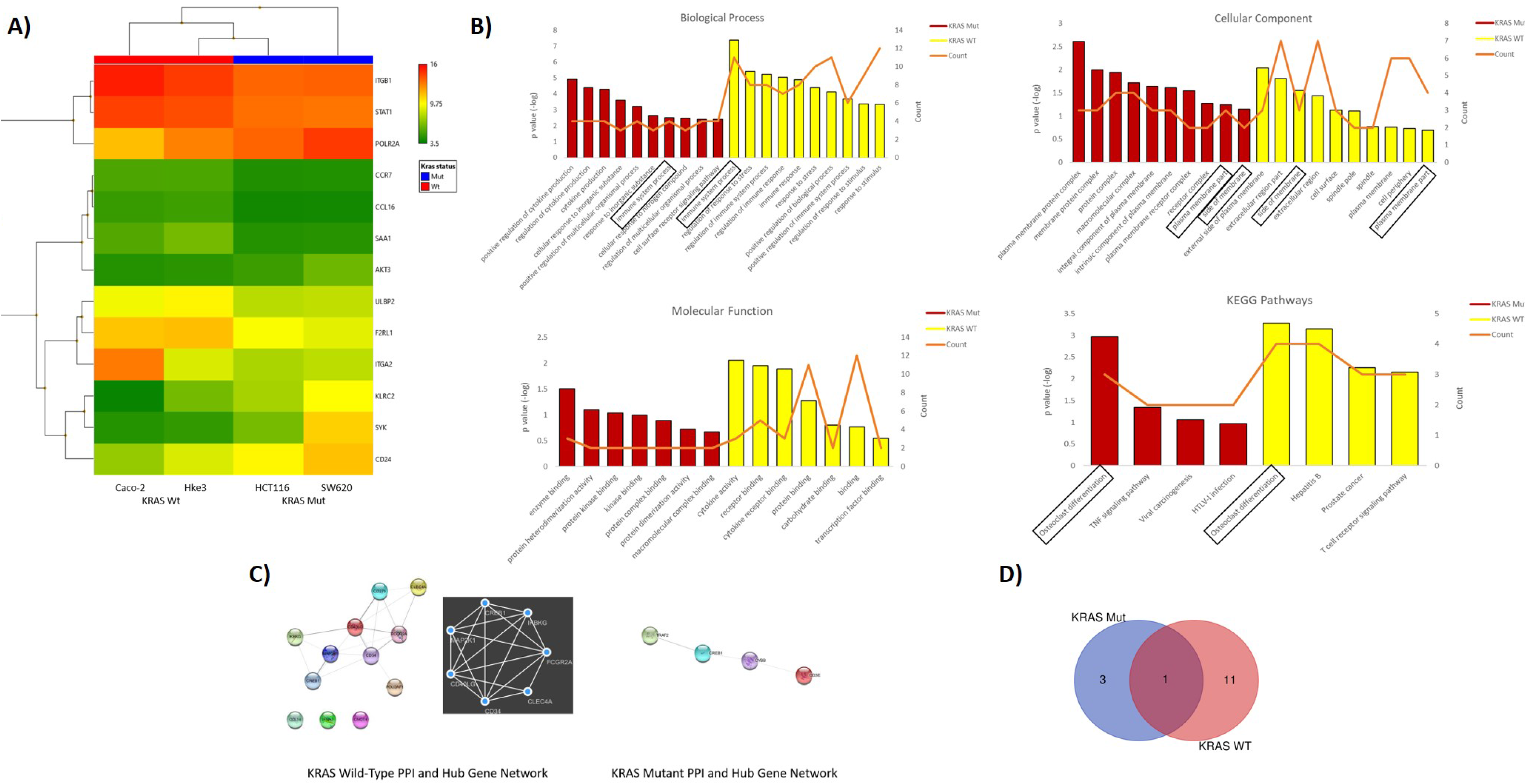
Differential expression of global immune response genes under reovirus infection in CRC cell lines categorized on *KRAS* status. (A) Hierarchical-clustering profile across differentially expressed genes (DEGs) participating in immune pathways upon treatment with reovirus in *KRAS*^Mut^ and *KRAS*^Wt^ groups. RMA normalized expression values for the 13 genes were used to generate a heat-map in the TAC software. The colors indicate the expression value relative to the median expression value per gene in the dataset. Red indicates upregulation relative to median value and green indicates downregulation relative to the median value. (B) GO term enrichment and KEGG pathway analyses of DEGs using DAVID. Differentially expressed probes between *KRAS*^Mut^ and *KRAS*^Wt^ cell lines treated with reovirus were selected based on meeting the criteria of p ≤ 0.05, and divided into positive and negative fold changes lists and used to determine enrichment. (C) Protein interaction analysis using STRING. The different colored nodes represent different genes. Different edges (lines) show the interplay between genes. Edge strength is shown by thickness of the line, the thicker the line, the stronger the interaction. Hub networks made using MCODE demonstrate the genes within the PPI that were screened to display the most highly upregulated. The genes not included in the hub networks did not have substantial interaction amongst the other genes in the list, therefore they were not included. (D) Venn diagram created using the TAC software. It illustrates the clear point that between the KRAS subgroups, CREB1 gene is the only commonality out of 16 genes, 4 in the *KRAS*^Mut^ and 12 genes in the *KRAS*^Wt^ groups.

**Supplementary Fig. 2.**
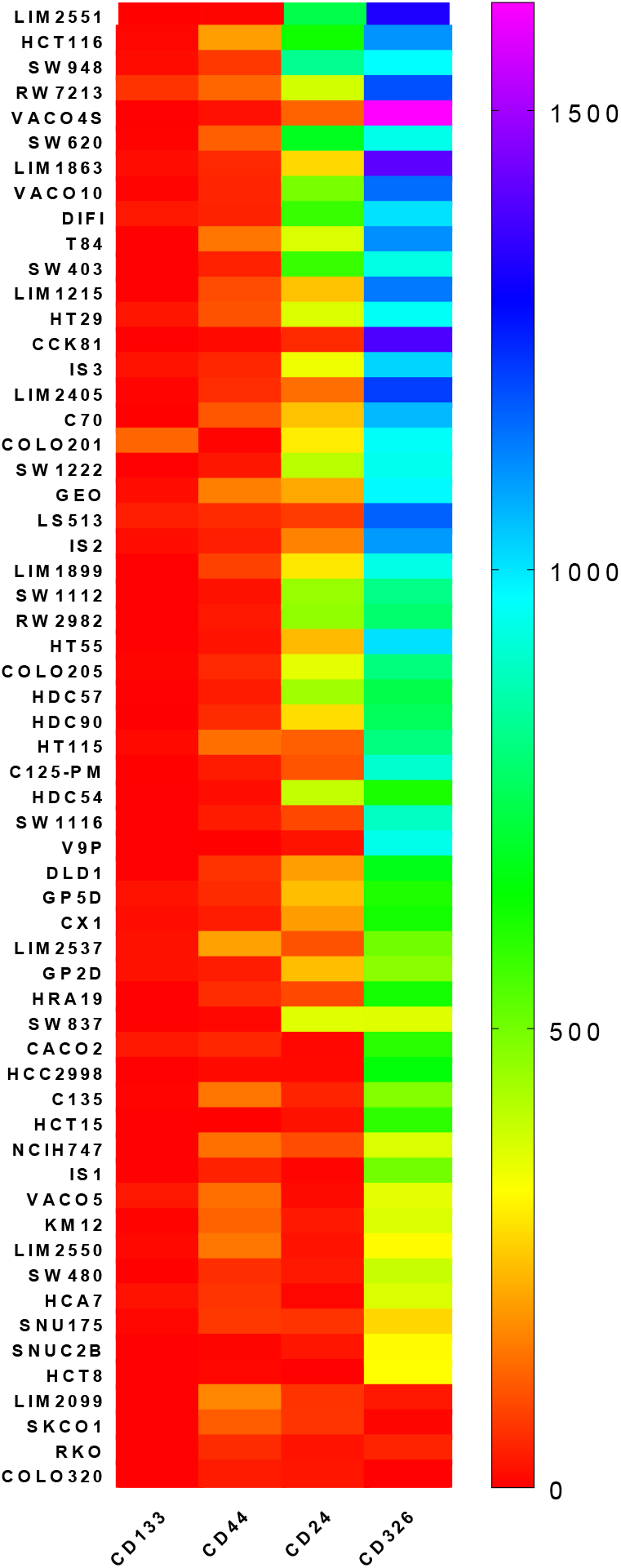
Classification of 59 CRC cell lines based on expression of stemness markers. Heat-map of stemness and epithelial cell markers’ expression among CRC cell lines studied using RNA-Seq. Transcript levels are given as reads per kilobase per million mapped (RPKM) values.

**Supplementary Fig. 3.**
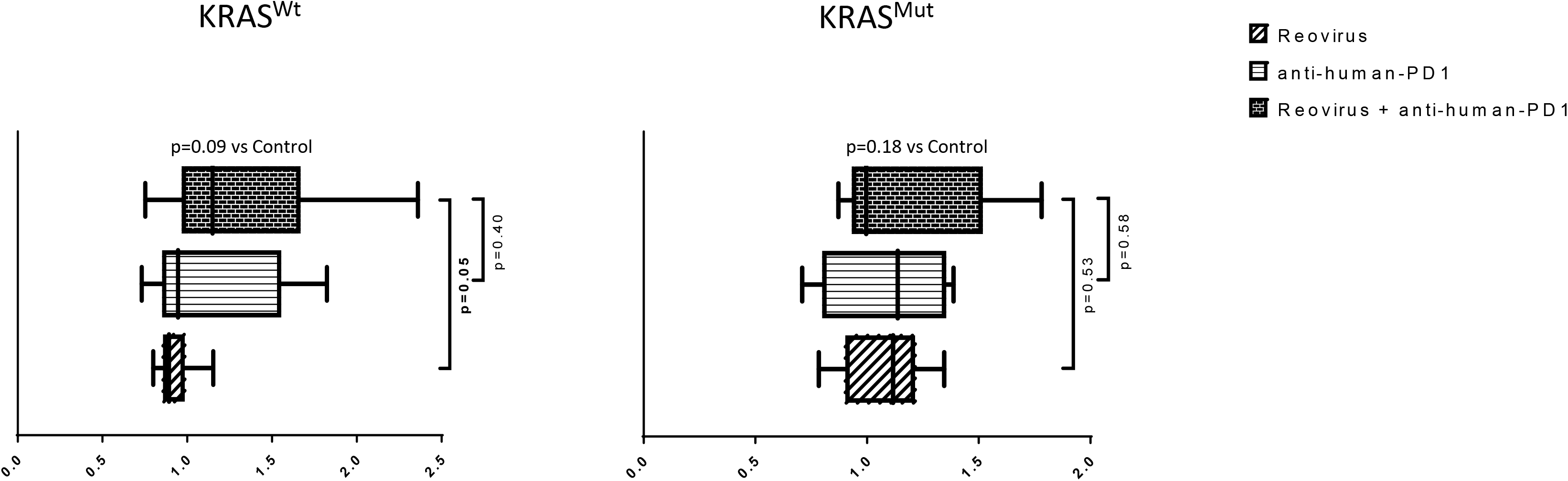
Levels of cell death among *KRAS*^Wt^ and *KRAS*^Mut^ human CRC cell lines co-cultured with human PBMC (*ex vivo*) and treated with reovirus, anti-human PD-1 and their combination. Trends in fold difference and significance are depicted on box and whisker plot. Combination treatment compared to placebo rendered no significance in either groups (p values in bold letters indicate significance).

**Supplementary Fig. 4.**
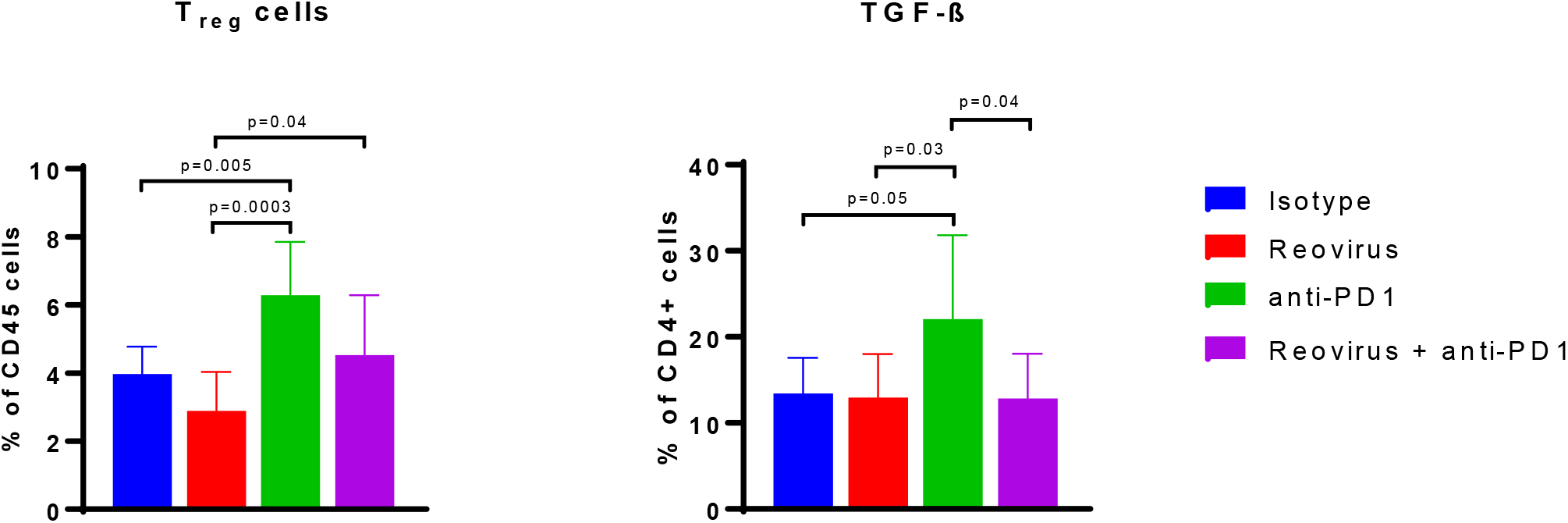
Alterations in Treg cell populations and TGF-β expression in CD4^+^ T cells upon treatment with reovirus and anti-PD-1 on CT26 tumors *in vivo*. While both single agents displayed significant role, combo treatment compared to control kept regulatory T cell levels unaltered. Interestingly enough, while reovirus treatment reduced, anti-PD-1 alone significantly increased Tregs in the TME of CT26. TGF-β, an immunosuppressive marker, by following the trends in Tregs was increased by single agent anti-PD-1 treatment. Combo treatment compared to control and single agent reovirus kept TGF-β expression unaltered.

**Supplementary Fig. 5.**
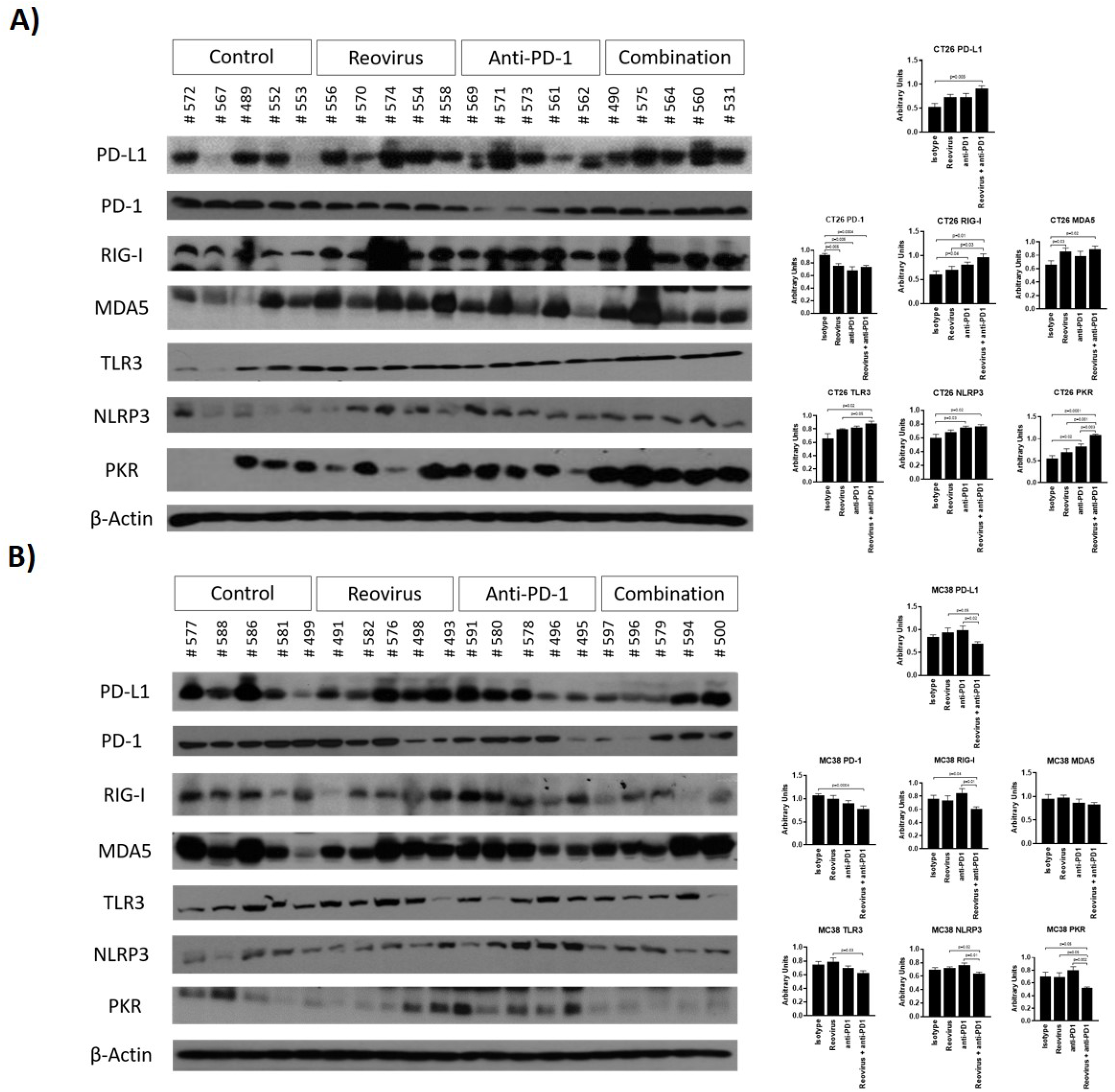
Protein level expressions of PD-L1, PD-1 and PRRs among individual tumor samples. Reovirus infection in CT26 upregulated most of the PRRs studied and either unaltered or decreased in expression, except for MDA5, in MC38. Cytoplasmic PRRs, RIG-I and MDA5, significantly increased upon combination treatment in CT26, whereas decreased in MC38. Combo treatment significantly increased membrane bound TLR3 in CT26, whereas reduced (p=NS) in expression in MC38. NLRP3, which is an inflammasome activation mediator, significantly increased by protein expression in CT26 TME, whereas mostly unaltered under any treatment conditions in MC38. dsRNA-sensing PRR, PKR, significantly increased by combo in CT26, however, reduced in expression in MC38. Densitometry quantifications (n=5) are given on bar charts.

### Supplementary Movie

**Reovirus and anti-human PD-1 (nivolumab) combinatorial treatment increases cell death among MSS compared to MSI cell line in ex vivo co-culture system**. Cytotoxicity of reovirus and anti-human PD-1 towards cell lines was monitored by time-lapse live-cell imaging (Cytotox Red uptake and release) over time, and found its significant increase among HT29 (MSS) cells. Combo treatment was comparatively ineffective in cell killing in LIM2405 (MSI). Statistical analysis is shown for the 4 hour and 9 hour co-incubation time points. Data represent at least 3 independent experiments (n=12). Mean values ± SEM are calculated.

(HT29 and LIM2405 Movies.mp4 in Incucyte folder)

